# Red fluorescent labeling of myelin by membrane-targeted tdTomato in transgenic mouse lines

**DOI:** 10.64898/2026.04.17.718425

**Authors:** Anja Reinert, Ulrike Winkler, Sandra Goebbels, Lina Komarek, Wiebke Möbius, Henri Seraf Zanker, Robert Fledrich, Ruth M. Stassart, Petra G. Hirrlinger, Klaus-Armin Nave, Hauke B. Werner, Aiman S. Saab, Johannes Hirrlinger

**Author notes:** **Correspondence** Prof. Dr. Johannes Hirrlinger. **Author contributions** Conceptualization: ASS, JH; Data curation: AR, JH; Formal analysis: AR; Funding acquisition: ASS, KAN; Investigation: AR, UW, SG, LK, HSZ, RF, RMS; Methodology: AR, UW, SG, RF, RMS, PGH, KAN, ASS, JH, WM; Project administration: JH; Software: AR; Supervision: JH; Validation: JH; Visualization: AR; Writing – original draft: AR, HBW, JH; Writing – review & editing: all authors.

## Abstract

Myelin is a highly complex membranous structure wrapped around axons by oligodendrocytes or Schwann cells in the central and peripheral nervous system, respectively. Fluorescent labeling is widely used to study the structure and dynamics of myelin. Combining structural with functional imaging requires labeling of myelin with red fluorescence, as many functional sensors, including Ca^2+^ indicators and genetically encoded metabolite sensors, fluoresce in the green spectral range. However, in vivo tools enabling red fluorescent labeling of myelinating cells and their myelin sheaths remain limited. Here, we generated a set of seven transgenic mouse lines expressing a membrane-targeted variant of the red fluorescent protein tdTomato in myelinating oligodendrocytes and Schwann cells throughout the nervous system. The mouse lines provide a variety of expression patterns ranging from wide-spread labeling of myelin to a rather sparse expression, the latter enabling visualization of individual oligodendrocytes and their associated myelin sheaths. In the peripheral nervous system, the pattern of fluorescence in sciatic nerves indicates predominant localization of tdTomato to non-compact myelin compartments including the inner and outer tongues, paranodal loops and Schmidt-Lanterman incisures. In summary, our work provides a set of novel mouse lines with myelin labeled by red fluorescence, which are compatible with diverse imaging modalities in the green spectral range enabling integrated structural and functional imaging.

**Main Points:**

- Transgenic mouse lines expressing membrane-targeted tdTomato in myelin enable imaging of myelin in the red spectral range
- Distinct expression patterns range from wide-spread labeling to sparse single-cell resolution, supporting diverse imaging applications

## Introduction

Myelin sheaths facilitate rapid, saltatory impulse propagation along the axons they myelinate (Tasaki, 1939; Cohen et al., 2020) and thus enable normal motor, sensory, and cognitive functions of the nervous system (Seidl et al., 2014; Bercury and Macklin, 2015). During development, myelinating cells follow an intrinsic program of proliferation, migration, and differentiation that is controlled by exogenous cues, including axonal surface molecules (Ahrendsen and Macklin, 2013; Mensch et al., 2015; Salzer et al., 2024). The formation of a myelin sheath includes establishment and stabilization of an initial contact between an axon and the tip of a cellular process of the prospective myelinating cell and the radial and longitudinal expansion of the myelin membrane (Hughes et al., 2013; Snaidero et al., 2014; Salzer et al., 2024). By electron microscopy, mature cross-sectioned myelin sheaths consisting of multiple membrane layers appear morphologically almost alike when comparing central and peripheral nervous system (CNS, PNS). However, oligodendrocytes and Schwann cells, the respective myelinating cells, differ in developmental origin, number of myelinated axonal segments per myelinating cell, myelin periodicity, transcription factor network, and myelin protein composition (Nave and Werner, 2021). Although mature myelin membranes are largely compacted, the adaxonal (innermost) and abaxonal (outermost) layers provide non-compacted, metabolically active myelin subcompartments, which are connected via cytosolic channels across the otherwise compacted myelin layers, including at the paranodes and in the internodal segment (Arroyo and Scherer, 2000; Edgar et al., 2021; Chapple et al., 2024); in the PNS the latter are termed Schmidt-Lanterman incisures. In the CNS, oligodendrocytes display regional heterogeneity with respect to morphological and electrical properties, transcriptional profiles, and myelin protein composition (Bakiri et al., 2011; Chong et al., 2012; Marques et al., 2016; Seeker et al., 2023; Siems et al., 2025). The capacity of oligodendrocytes to react to local cues thus appears as prerequisite for refining their functions. Whether an axonal segment is myelinated or not remains critical for its impulse propagation properties beyond development. Indeed, myelination can be triggered in adults by functional activity (Bengtsson et al., 2005; Scholz et al., 2009; Liu et al., 2012; Sampaio-Baptista et al., 2013) as a mechanism of plasticity. On the other hand, dys- and demyelination or the transition of oligodendrocytes to a disease-associated state are key facets of the pathophysiology observed in myelin-related disorders (Falcão et al., 2018; Wolf et al., 2021; Depp et al., 2023; Stadelmann et al., 2025).

Current myelin biology relies critically on tools that allow measuring the presence and morphological properties of myelin sheaths in rodent models (Chong et al., 2012; Dereddi et al., 2026), as well as tools to study the molecular (Eichel et al., 2020; Cai et al., 2025) and metabolic (Looser et al., 2018; Trevisiol et al., 2020; Looser et al., 2024) interactions between axons and their myelin sheaths. Several mouse lines have been previously generated that transgenically label oligodendrocytes and/or myelin. For instance, cytosolic Enhanced Green Fluorescent Protein (EGFP) was expressed under control of the proteolipid protein (*Plp1)*-promoter (Mallon et al., 2002) or the myelin-associated oligodendrocyte basic protein (*Mobp*)-promoter (Gong et al., 2003; Hughes et al., 2018). Mouse lines with EGFP localized to the plasma membrane via the CAAX membrane anchor domain (Hancock et al., 1989) were generated using the myelin basic protein (*Mbp)*-promoter (Chong et al., 2012) or the cyclic nucleotide phosphodiesterase (*Cnp)*-promoter (Deng et al., 2014). While these mouse lines provide labeling of myelinating cells and their myelin sheaths, the fluorescence of EGFP is in the green spectral range thereby spectrally overlapping with many functional sensors including green fluorescent calcium sensors (Zhang and Looger, 2024) and most of the genetically encoded sensors for metabolites (Koveal et al., 2020; San Martín et al., 2022). This obstacle can be overcome by imaging myelin with label-free techniques (Schain et al., 2014; Craig et al., 2024; Kamen et al., 2025) or by labeling of oligodendrocytes and myelin with a red fluorophore. Indeed, a mouse line expressing the red fluorescent protein DsRed under control of the *Plp1* promoter has been established (Hirrlinger et al., 2005). However, in this mouse line the fluorescent protein mainly localizes to the cytosol, thus providing labeling of oligodendrocyte cell bodies but only limited signal in myelin sheaths (Hirrlinger et al., 2005). Alternatively, red-fluorescent labeling of cytosolic channels in myelin has been achieved by combining *Plp1*-CreERT2 mice with tdTomato expression dependent on Cre-mediated DNA recombination (Edgar et al., 2021), a strategy which requires crossbreeding of two mouse lines and tamoxifen application. However, to the best of our knowledge, no single transgenic, Cre-independent mouse line currently provides membrane-targeted red fluorescence enabling robust visualization of myelinating cells and their myelin sheaths.

Overcoming this limitation, we here report a set of novel transgenic mouse lines that express the membrane-targeted bright red fluorescent protein tdTomato (Ex/Em= 554/581 nm) in myelinating cells driven by the *Cnp*-promoter. A variety of expression patterns was observed in the different lines, ranging from wide-spread to sparse oligodendroglial expression, with sparse lines enabling visualization of individual oligodendrocytes and their associated myelin sheaths. These mouse lines provide a valuable tool for myelin research as they will allow to combine red fluorescent morphological labeling of myelin with imaging of green fluorescent functional sensors.

## Material & Methods

### Ethics statement

All animal experiments were performed in accordance with the guidelines outlined by the European Communities Council Directive (2010/63/EU), the German Protection of Animals Act (TSchG), as well as guidelines of the Max Planck Society and received approval from the animal welfare office of the Max Planck Institute for Multidisciplinary Science, as well as from the local governmental authorities (Niedersächsisches Landesamt für Verbraucherschutz und Lebensmittelsicherheit; LAVES). Generation of transgenic mice was performed under license 33.19-42502-04-21/3660 issued by LAVES.

### Design of the transgene construct and generation of transgenic mice

To target a membrane-tagged version of the red fluorescent protein tdTomato to the myelin membrane, the transgene consisted of a 3919 bp long part of the promoter of 2’-3’-cyclic nucleotide 3’-phosphodiesterase (*Cnp*) (for full sequences see Suppl. Fig. 1), followed by the open reading frame of a codon-optimized variant of tdTomato attached to the CAAX membrane anchor sequence, which has been previously used to target EGFP to myelin membranes (Chong et al., 2012; Deng et al., 2014). On the 3’-end of the transgene, a late SV40 polyadenylation site was inserted (Suppl. Fig. 1). The DNA construct was synthesized by Vectorbuilder (Chicago, IL, USA) and obtained as microinjection-ready transgenic vector DNA preparation.

Pronuclear microinjection into oocytes of C57BL/6N mice was performed by the Laboratory for Transgenic Technologies of the Max Planck Institute for Multidisciplinary Sciences, Göttingen, Germany. Nine transgenic mice were born from the pronuclear microinjection, from which seven transgenic lines could be established (designated as lines A, C, D, E, F, H, I). Mice were maintained on the C57BL/6N genetic background. These mouse lines have been registered at Mouse Genome Informatics MGI as C57BL/6N-Tg(Cnp-tdTomato*)<Y>Jhi with <Y> = A, C, D, E, F, H, I for the different lines (Table 1). Within this MS, the nickname CNPMTO-<Y> (for *Cnp*-membrane-td<u>To</u>mato) is used.

**Table 1:** Official names and Mouse Genome Informatics registration numbers.

|  | allele |  | mouse line |  |
| --- | --- | --- | --- | --- |
| Line | name | MGI | name | MGI |
| A | Tg(Cnp-tdTomato*)AJhi | 8278415 | C57BL/6N-Tg(Cnp-tdTomato*)AJhi/Jhi | 8278537 |
| C | Tg(Cnp-tdTomato*)CJhi | 8278530 | C57BL/6N-Tg(Cnp-tdTomato*)CJhi/Jhi | 8278545 |
| D | Tg(Cnp-tdTomato*)DJhi | 8278531 | C57BL/6N-Tg(Cnp-tdTomato*)DJhi/Jhi | 8278546 |
| E | Tg(Cnp-tdTomato*)EJhi | 8278532 | C57BL/6N-Tg(Cnp-tdTomato*)EJhi/Jhi | 8278547 |
| F | Tg(Cnp-tdTomato*)FJhi | 8278533 | C57BL/6N-Tg(Cnp-tdTomato*)FJhi/Jhi | 8278548 |
| H | Tg(Cnp-tdTomato*)HJhi | 8278534 | C57BL/6N-Tg(Cnp-tdTomato*)HJhi/Jhi | 8278549 |
| I | Tg(Cnp-tdTomato*)IJhi | 8278535 | C57BL/6N-Tg(Cnp-tdTomato*)IJhi/Jhi | 8278550 |

### Genotyping

Mice were genotyped by routine genomic PCR using the following primers: forward: 5’-GAG CAG TAT GAA CGG AGC GA-3’ / reverse 5’-GGA TAC AGC CGT TCA GTG CT-3’ yielding a product of 548 bp. As an internal control, the following primers were included in the PCR, which amplify a 721 bp fragment of the SLC1A3 (GLAST) gene: forward 5’-GAG GCA CTT GGC TAG GCT CTG AGG A-3’ / reverse 5’-GAG GAG ATC CTG ACC GAT CAG TTG G-3’.

### Expression analysis of tdTomato

The expression of tdTomato was analyzed in adult mice in an age range between 54 days and 327 days (mean age 143 days, n=44). Mice were transcardially perfused with 4% paraformaldehyde (PFA) in phosphate buffer saline (PBS, pH 7.4). Brains, optic nerves and sciatic nerves were dissected and post-fixed in 4% PFA/PBS at 4°C for 24 h.

#### Brain section immunolabeling

Sagittal brain sections of 40 µm thickness were cut on a vibratome. Free-floating slices were permeabilized in 0.4% Triton X-100/PBS for 30 min and blocked with 4% fetal calf serum (FCS) in 0.2% Triton X-100/PBS for 1 h. Sections were immunohistochemically stained either for MBP (rat anti-MBP, AbD Serotec #MCA409S) or CNP (guinea pig anti-CNP, Synaptic Systems #355004). Each primary antibody was diluted 1:500 in primary antibody solution (1% FCS, 0.05% Triton X-100/PBS) in which the sections were incubated overnight at 4°C. After washing, slices were incubated in secondary antibody solution (1.5% FCS/PBS) for 2 h at RT containing one of the secondary antibodies (diluted 1:500): donkey anti-rat Cy5 (Jackson Immuno Research #712-175-153), donkey anti-rat DyLight755 (Invitrogen #SA5-10031), donkey anti-guinea pig Cy5 (Jackson Immuno Research #706-175-148) or goat anti-guinea pig DyLight755 (Invitrogen #SA5-10099). Sections were washed in PBS, mounted on SuperFrost slides (Thermo Fisher Scientific, USA) and embedded in Fluoromount-G (Invitrogen), which includes DAPI as nuclear stain.

#### Colocalization analysis on whole-section scans

Whole sagittal brain section scans were acquired using a Zeiss AxioScan.Z1 microscope (Zeiss Microscopy GmbH, Jena, Germany). Images were taken with a 20× objective (NA 0.8) at a single focal plane with a lateral resolution of 0.227 µm per pixel (x–y) and 16-bit. LED modules for excitation were 385 nm (DAPI, automatic focus), 567 nm (tdTomato) and 735 nm (DyLight755). Microscopic settings were identical for imaging of all brain slices. For presentation of fluorescence images of whole-section scans (Fig. 1) images were binned 2 × 2 with a final lateral resolution of 0.454 µm per pixel using ZEN 3.1 (Zeiss Microscopy GmbH, Jena, Germany). Grayscale and pseudocolored (LUTs, 16-colors) 16-bit images were generated with Fiji (National Institutes of Health, USA; Schindelin et al., 2012).

**Fig. 1:**
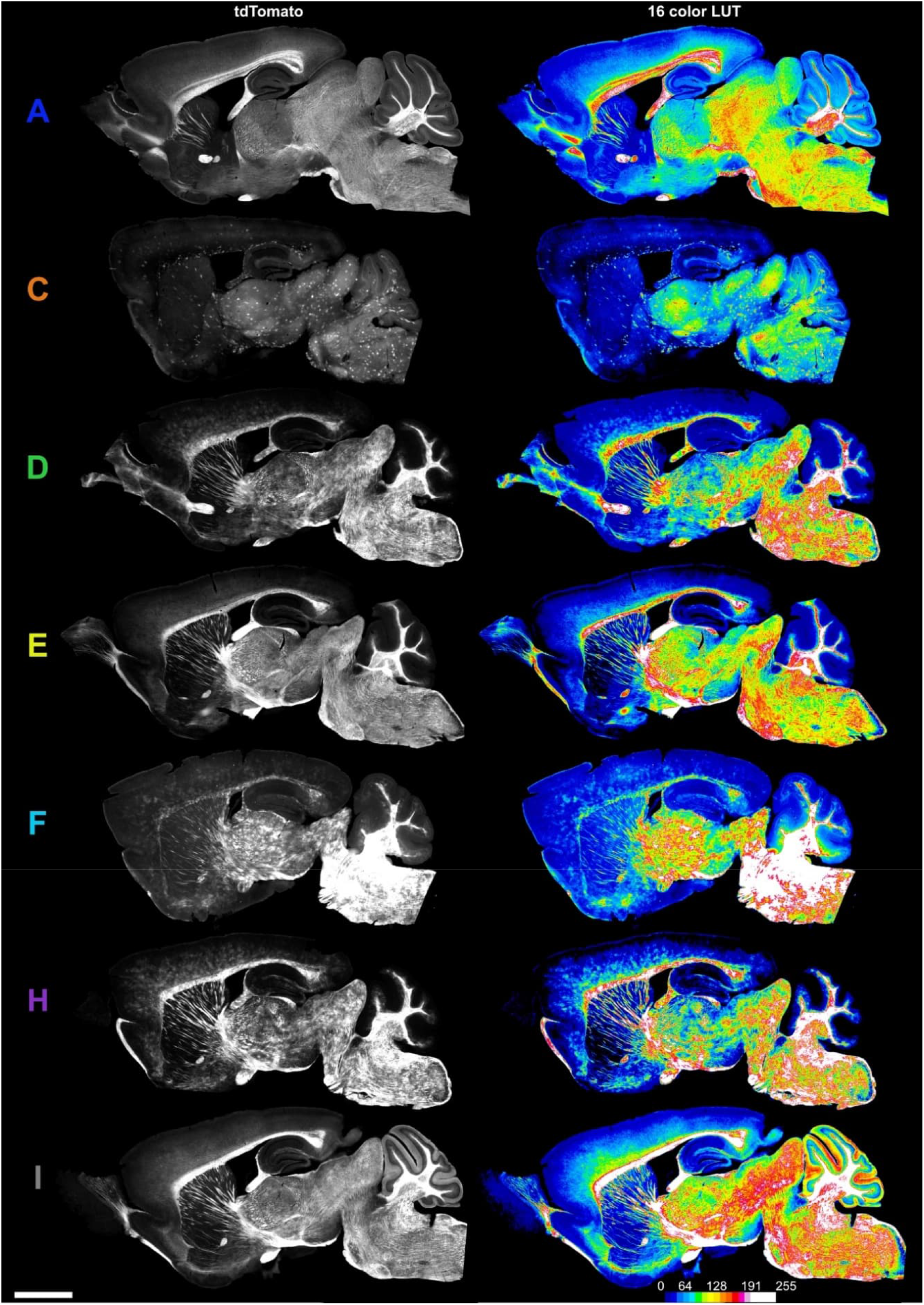
tdTomato fluorescence in the newly generated CNPMTO mouse lines. Grayscale (left) and pseudo-color (right) fluorescence images show the overall expression pattern of tdTomato in sagittal whole-brain sections of the seven transgenic mouse lines (CNPMTO-A, C, D, E, F, H, I). The color bar indicates fluorescence intensity (a.u.). The scale bar corresponds to 2 mm and applies to all panels. tdTomato shows a wide-spread (CNPMTO-A, E, I), patchy (CNPMTO-D, F, H) or sparse (CNPMTO-C) distribution. High-resolution images are available at Zenodo at https://doi.org/10.5281/zenodo.19592043.

Colocalization of transgenically expressed tdTomato fluorescence and MBP immunoreactivity was quantified using the BIOP JACoP plugin in Fiji (https://github.com/fabricecordelieres/IJ-Plugin_JACoP; Bolte and Cordelières, 2006). Prior to colocalization analysis, images underwent the following processing steps using ZEN 3.1: to reduce memory usage during image analysis, images were downsampled in x–y plane using 4 × 4 binning with a final lateral resolution of 0.98 µm per pixel. Images were subsequently deblurred (strengths: 0.9, radius: 15, sharpness: 0.0). The channels (tdTomato, MBP) of the processed images were separately exported as uncompressed tiff files. The two single channel files were transformed to 8-bit images with Fiji. To exclude background fluorescence each image was automatically intensity thresholded, and the two channels subsequently merged into a composite image. ROIs were manually set for the whole slice (section-wide), cortex (motor cortex region layer I-VI), corpus callosum, hippocampus, thalamus, midbrain, medulla and cerebellum. Composite images were analyzed with BIOP JACoP. As colocalization parameters the tdTomato channel was defined as channel A and the MBP channel as channel B. The threshold for both channels was set to Mean. This thresholding approach was verified to provide the highest precision, accurately detecting delicate myelin fibers without over-thresholding regions of dense signal. The fraction of MBP-positive fibers expressing tdTomato was quantified as area double positive for both tdTomato and MBP (area_tdTomato+MBP_) divided by the area stained for MBP (area_MBP_) both reported by BIOP JACoP. Diagrams were generated with GraphPad Prism 10 (GraphPad Software, San Diego, USA).

#### Confocal imaging of fixed brain sections

Confocal images of fixed brain sections were taken with the LSM 880 (Zeiss Microscopy GmbH, Jena, Germany) using a 63× water immersion objective (NA 1.2). Lasers with an excitation wavelength of 561 nm (tdTomato) and 633 nm (Cy5) were used. The confocal pinhole was set to 0.9 airy units (AU). Stacks along the *z*-axis were taken at distances of 1.5 μm. Average intensity projection images were generated and auto-adjusted for brightness and contrast with Fiji.

#### Confocal imaging of fixed optic and sciatic nerves

Optic and sciatic nerves were whole-mounted (Fluoromount-G, Invitrogen) on SuperFrost slides. For cross-sections, sciatic nerves were cryoprotected in sucrose, embedded in Tissue-Tek O.C.T. compound (Sakura Finetek), cross-sectioned on a cryostat (10 µm) and mounted (Aqua-Poly/Mount, Polyscience) on SuperFrost slides. Confocal images were taken with the Zeiss LSM 880 using the Airyscan mode and a 20× objective lens (NA 0.8). Stacks along the *z*-axis were taken at distances of 0.5 μm. Average intensity projection images were generated and adjusted in brightness and contrast with Fiji.

#### Teased sciatic nerve fibers

Fixed sciatic nerves of line CNPMTO-A (age 165 days) were microdissected in PBS under a stereomicroscope. The epi- and perineurium were removed with fine forceps and small fiber bundles were transferred into a drop of PBS on a SuperFrost slide. By using needles individual fibers were subsequently teased from the bundles. From selected nerve fibers the endoneurium was removed using fine forceps. Excess liquid was removed and nerve fibers were air-dried and mounted (Fluoromount-G, Invitrogen). Epifluorescence images were taken with an Axio Observer.Z1 (Zeiss Microscopy GmbH, Jena, Germany) using a 40× C-Apochromat water immersion objective (NA 1.2).

#### Immuno-transmission electron microscopy of optic nerves

Immunogold-labeling of tdTomato in cryo-sectioned cross-sections of optic nerves and subsequent TEM was performed as previously described (Weil et al., 2019). In brief, mice were perfused with 4%PFA/PBS and the brain with the attached optic nerves was isolated and post-fixed in 4% PFA, 0.2% glutaraldehyde, 0.5% NaCl in PBS overnight at 4°C. For storage the fixative was replaced by 1% PFA/PBS. For ultrathin cryosectioning, nerve pieces were embedded in small blocks of 10% gelatin and infiltrated overnight with 2.3 M sucrose in PBS. The blocks were mounted on aluminum pins for ultramicrotomy and were frozen in liquid nitrogen. Sections were cut on a UC6 cryo-ultramicrotome (Leica Microsystems, Wetzlar, Germany) using a 35° cryo-immuno-diamond knife (Diatome, Biel, Switzerland). For immunolabeling, sections were incubated with an antibody directed against RFP (# 600-401-379; polyclonal rabbit, Rockland Inc., 1:100) followed by protein A-gold (10 nm; Microscopy Center, Department of Cell Biology, University Medical Center Utrecht, The Netherlands). Sections were examined on a Zeiss LEO EM912AB (Zeiss Microscopy GmbH, Oberkochen, Germany) and digital images were acquired with a 2048 × 2048 CCD camera (TRS, Moorenweis, Germany).

## Results

To enable red fluorescent labeling of myelin, we generated transgenic mice expressing membrane-bound tdTomato under the control of the *Cnp*-promoter. Pronucleus injection of the transgene construct resulted in seven transgenic mouse lines expressing the fluorescent protein in myelinating cells as well as within myelin (see Table 1 for official names and MGI registration numbers). Within this MS, the nickname CNPMTO-<Y> (for *Cnp*-membrane-td<u>To</u>mato) is used, with <Y> = A, C, D, E, F, H, I for the different lines. Analysis of the expression pattern revealed a wide-spread expression of tdTomato in myelin in the brain in all mouse lines (Fig. 1; high-resolution images are available at Zenodo at https://doi.org/10.5281/zenodo.19592043). However, the detailed expression pattern differed between lines (Fig. 1, 2). In CNPMTO-A, -E, and -I wide-spread tdTomato fluorescence was observed in many, rather equally distributed oligodendrocytes with strong labeling of myelin structures. Lines CNPMTO-D, -F, and -H presented with a patchier expression pattern in which clusters of oligodendrocytes express tdTomato, while other cells between these clusters are not fluorescent. Finally, line CNPMTO-C presents a sparse expression where only few cells and their myelin are labeled. This variability provides flexibility for different experimental applications, ranging from population-level visualization to single-cell resolution imaging.

**Fig. 2:**
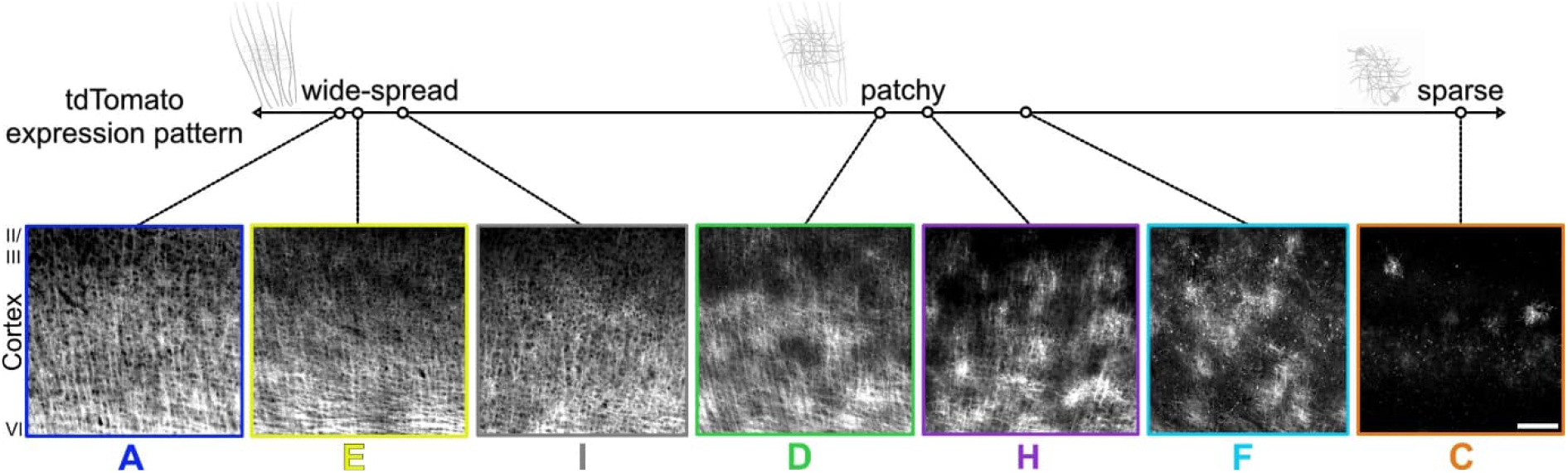
tdTomato expression patterns in CNPMTO lines are categorized as wide-spread, patchy, and sparse. Contrast-enhanced fluorescence micrographs of the cortex of each mouse line show the pattern of fluorescent myelin fibers including the different density of labeling. Line CNPMTO-A, E and I show wide-spread tdTomato expression in the majority of the myelinated fibers. Line CNPMTO-D, F and H show a patchy expression. In line CNPMTO-C only single oligodendrocytes are tdTomato-positive. The scale bar in the rightmost panel corresponds to 100 µm and applies to all panels.

To quantify the fraction of myelin labeled by expression of the membrane-bound tdTomato protein, myelin was stained with anti-MBP-antibodies and the MBP-immunopositive area was analyzed for tdTomato expression across whole sagittal brain slices as well as specific brain areas (Fig. 3). Consistent with the observed sparse tdTomato expression, the lowest labeling of myelin by tdTomato was observed in line CNPMTO-C across all brain regions. In contrast, lines CNPMTO-A, -E, and -I showed the highest labeling of myelin reaching between 60% and 80% of the MBP-immunopositive area across most brain regions. These values indicate that a substantial fraction of myelin is labeled in high-expression lines, while sparse lines provide selective labeling.

**Fig. 3:**
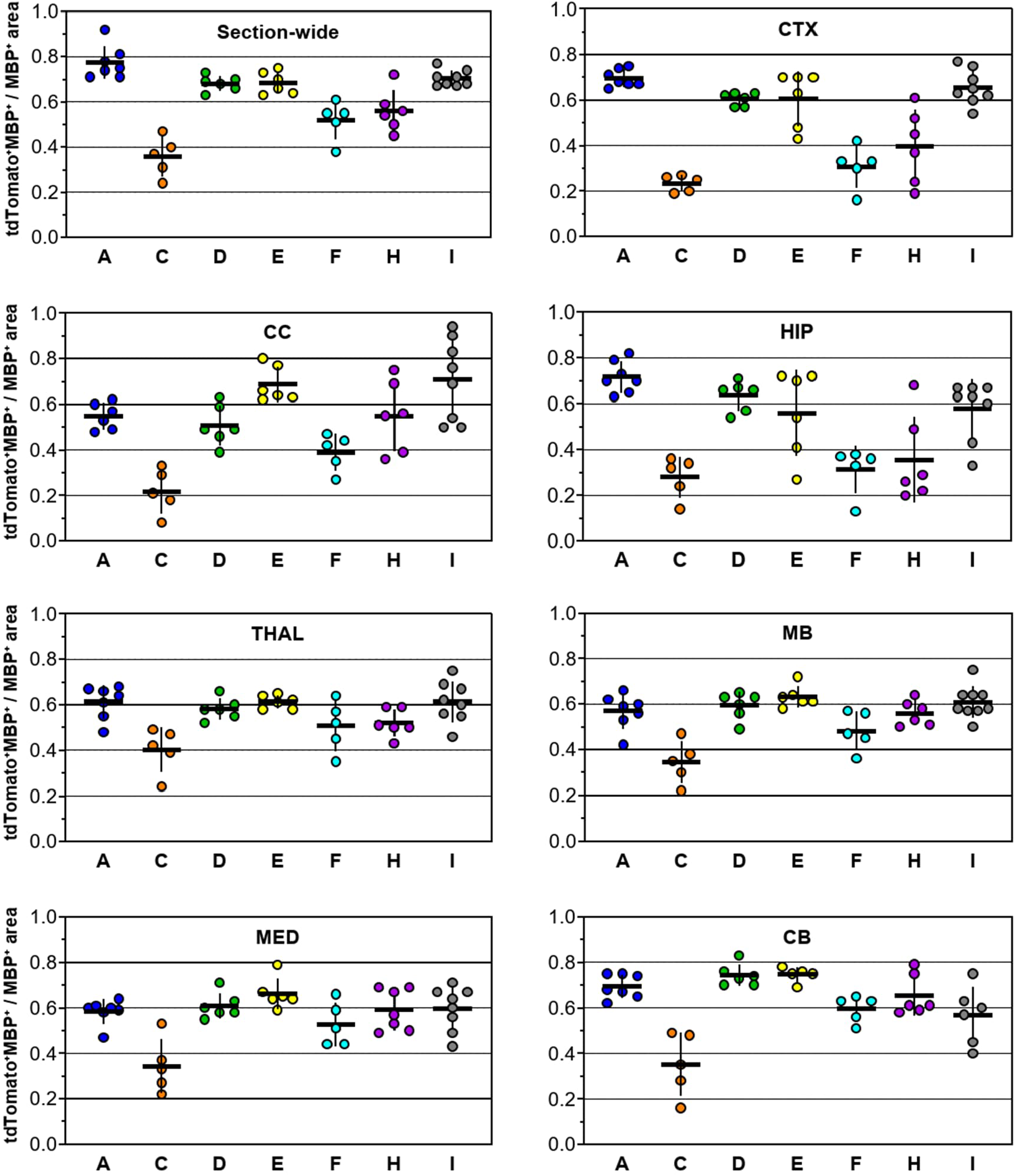
Quantification of the ratio of the MBP-immunopositive area labeled by tdTomato fluorescence in different brain regions. The ratio of tdTomato^+^MBP^+^ double-fluorescence positive area to total MBP^+^-immunopositive area is shown for the seven mouse lines (CNPMTO-A, C, D, E, F, H, I) analyzed for whole sagittal slices (‘Section-wide’) or the indicated brain regions. Each data point refers to one mouse (n=5-6 mice). The horizontal line indicates the mean, while the vertical line spans the standard error of the mean. CB: cerebellum; CC: corpus callosum; CTX: cortex; HIP: hippocampus; MB: midbrain; MED: medulla oblongata; THAL: thalamus.

Confocal imaging at higher magnification revealed localization of tdTomato within myelin along axons within different brain regions (Fig. 4). While in line CNPMTO-A a dense network of myelin sheaths was observed in gray and white matter regions, sparse labeling in line CNPMTO-C allowed to image single oligodendrocytes, their processes and associated myelin sheaths. In CNPMTO-H mice, an intermediate labeling pattern was observed. To further characterize the localization of the tdTomato fluorescence signal, co-immunolabeling with antibodies against either the oligodendrocyte marker CNP or the myelin marker MBP was performed (Fig. 5). In CNPMTO-H mice, tdTomato expression was frequently observed in the cell body of CNP-immunopositive oligodendrocytes, while in CNPMTO-C mice rather few CNP-immunopositive somata show red fluorescence (Fig. 5A). However, tdTomato fluorescence clearly labels myelin sheaths in both lines, which is also confirmed by MBP-co-immunolabeling (Fig. 5B). In many cases, myelin sheaths around the axon were clearly distinguishable, while the enwrapped axon itself remains unlabeled (Fig. 5B).

**Fig. 4:**
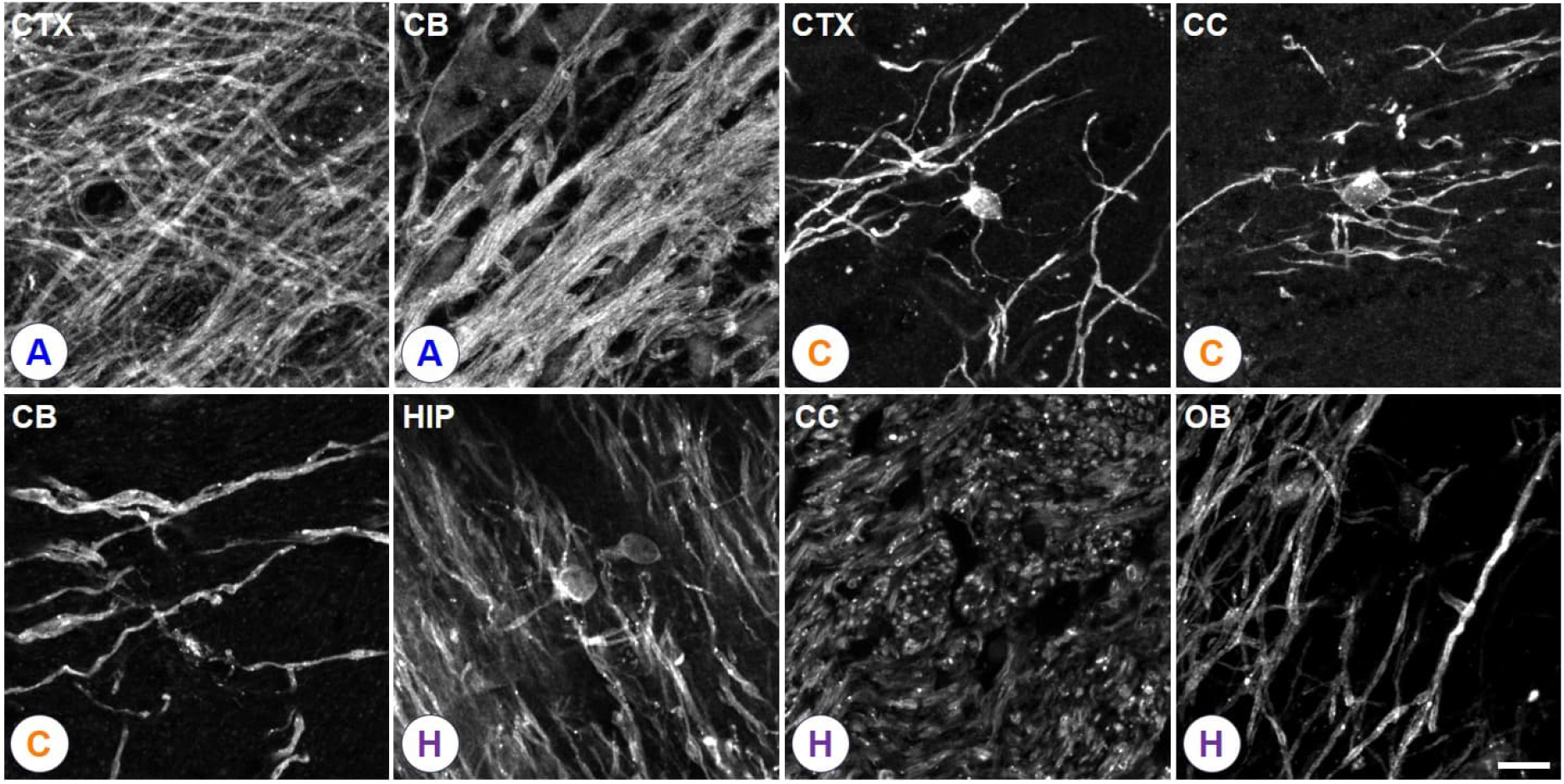
Examples of tdTomato fluorescence in oligodendrocytes and myelin in the brain. Shown are confocal images of gray and white matter regions of the mouse lines indicated in the circles with wide-spread (A), patchy (H) or sparse (C) expression. CB: cerebellum; CC: corpus callosum; CTX: cortex; HIP: hippocampus; OB: olfactory bulb. tdTomato expression highlights the soma and processes of oligodendrocytes and their myelin sheaths. The scale bar corresponds to 10 µm and applies to all panels.

**Fig. 5:**
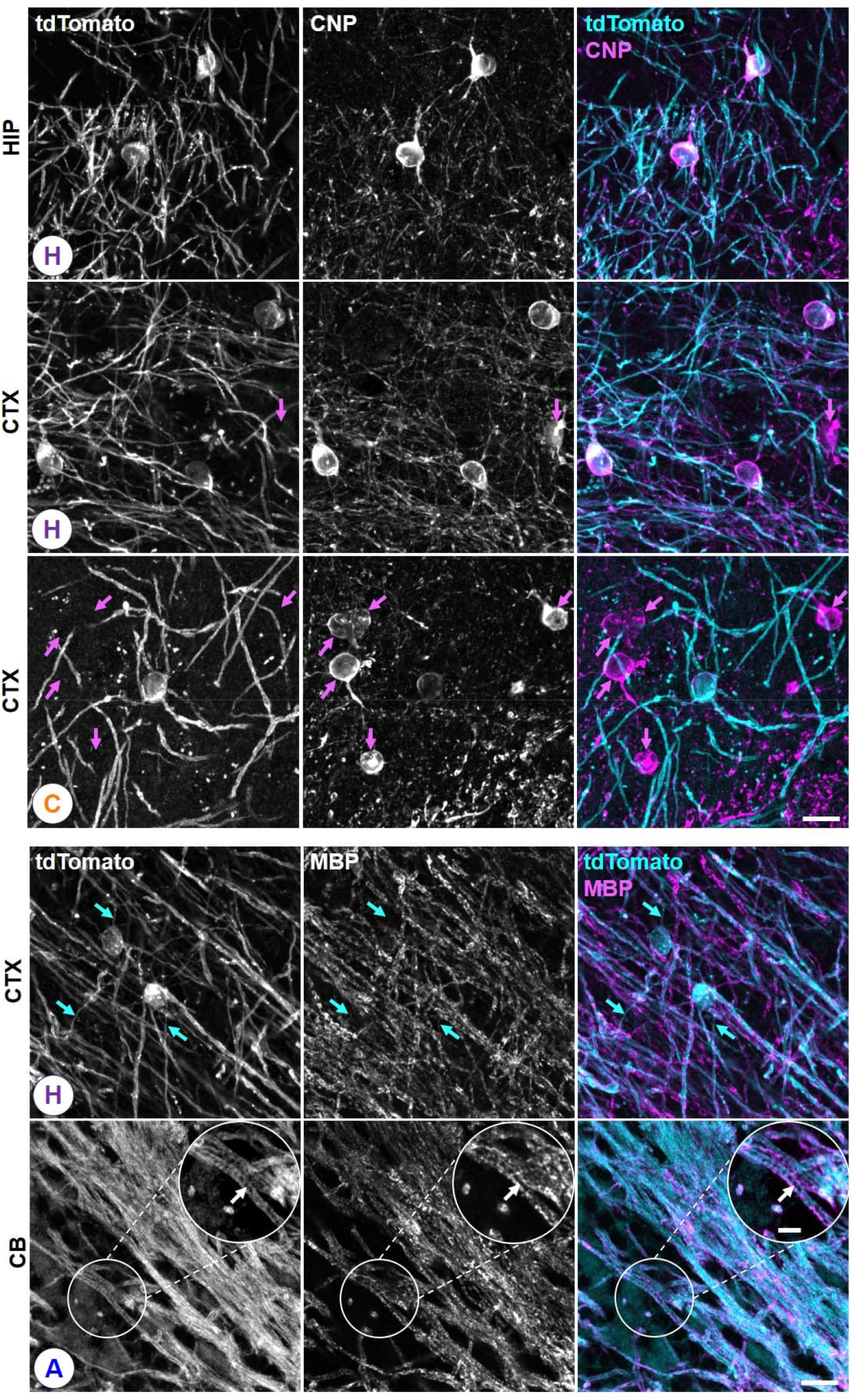
Immunolabeling of CNP and MBP in CNPMTO mice. Upper panels: Fluorescence micrographs visualizing that transgenically expressed tdTomato is localized in the processes and somata of oligodendrocytes which are marked by CNP immunolabeling. A fraction of CNP-positive oligodendrocytes does not express tdTomato (purple arrows), corresponding to the expression pattern as described in Fig. 2. Lower panels: Colocalization of MBP and tdTomato in myelin. The fluorescence signal is observed on both sides of the axon, while the enwrapped axon shows no fluorescence (white arrow in the inset). The cyan arrows highlight primary processes of oligodendrocytes connecting soma and myelin sheaths, which are labeled by tdTomato, but not stained for MBP. CB: cerebellum; CTX: cortex; HIP: hippocampus. The letters in the circles indicate the CNPMTO mouse line. Scale bar: 10 µm. Scale bar inset: 4 µm.

Optic nerves dissected from all seven transgenic lines were analyzed for tdTomato fluorescence in this CNS white matter tract (Fig. 6). Across all transgenic mouse lines, labeling of myelin was observed with the fluorescence outlining myelin sheaths around the axons. However, the number of tdTomato expressing oligodendrocytes in the optic nerve differed between the mouse lines following the pattern observed in the brain. Specifically, lines CNPMTO-A, -E, and -I appeared to label a large fraction of myelin, while only few oligodendrocytes express tdTomato in line CNPMTO-C, allowing to image single cells and their myelin sheaths (Fig. 6). Transmission electron microscopy (TEM) of immunogold-labeled optic nerve cross sections was used to localize tdTomato within the myelin in line CNPMTO-I (Fig. 6). Most of the immunogold-labeling was observed in the cytosolic compartment of the myelin sheath, while only few gold particles were localized in compact myelin.

**Fig. 6:**
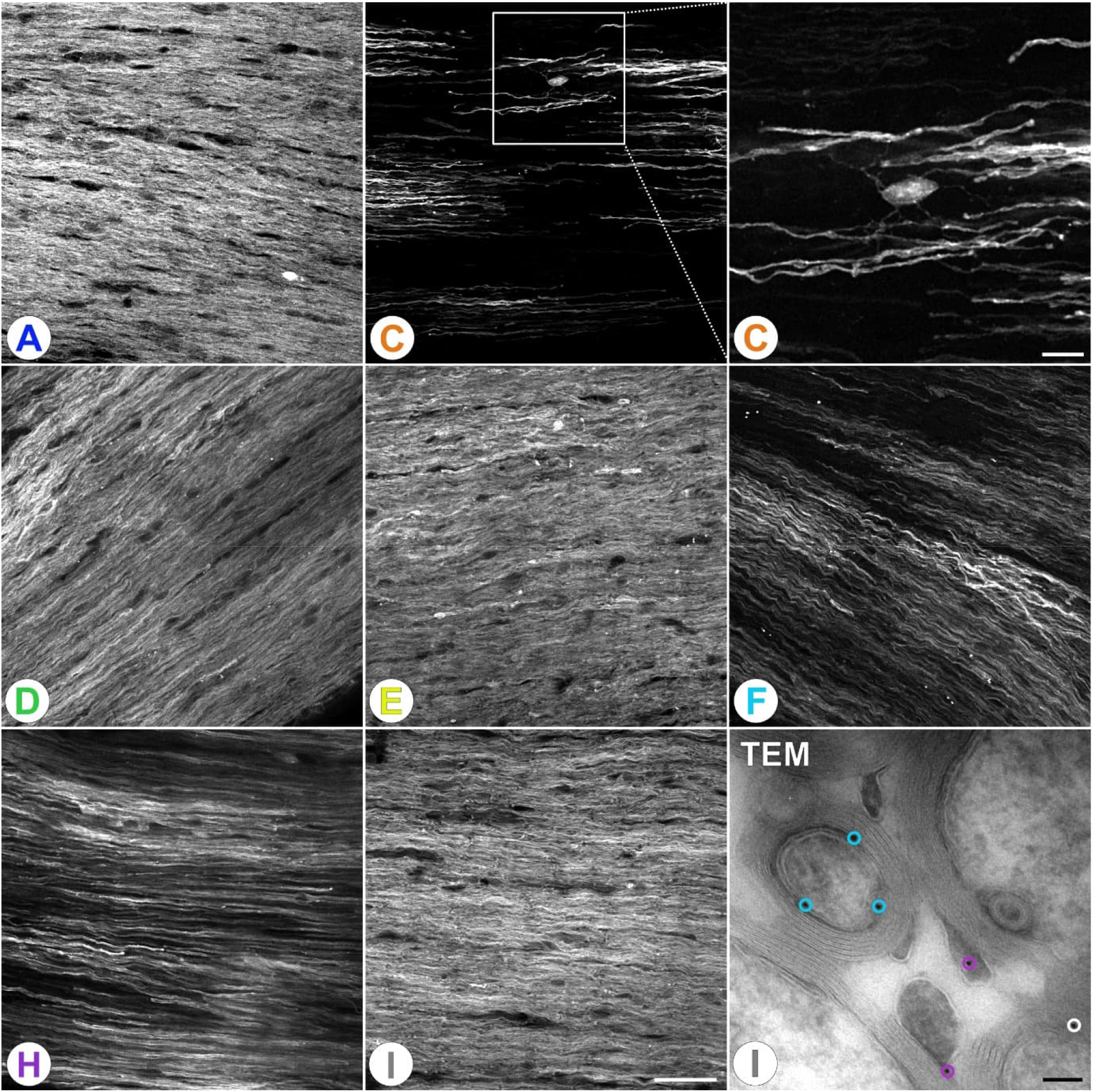
tdTomato expression in the optic nerves of the CNPMTO mouse lines. Confocal images of wholemount optic nerve preparations reveal line-dependent expression patterns ranging from wide-spread labeling of myelin to labeling of single oligodendrocytes and their associated myelin sheaths. TEM: Transmission electron micrograph of immunogold-labeled tdTomato protein in an ultrathin cross-section of the optic nerve of a CNPMTO-I mouse. tdTomato is associated with oligodendrocyte membranes and mainly localized in the inner (black gold particles with cyan circle) and outer (black gold particles with purple circles) cytoplasmic tongues and in electron-lucent cytoplasmic clefts interrupting the compact myelin sheath (black gold particles with white circle). The letters in the large white circles indicate the CNPMTO mouse line. Scale bar confocal images: 40 µm, inset 10 µm. Scale bar TEM image: 100 nm.

Finally, to assess the endogenous tdTomato expression within the PNS, sciatic nerves were analyzed as wholemounts, cross-sections and teased fibers. Confocal imaging visualized different expression patterns across the seven transgenic lines (Fig. 7) that exhibit broadly similar characteristics as in the CNS. A wide-spread expression of tdTomato in most fibers is present in CNPMTO-A, E and I, of which CNPMTO-E showed an overall weaker expression level. Lines CNPMTO-D and H present a patchy-like expression, with some bundles of fibers displaying a strong tdTomato fluorescent signal, while other fibers show weak or no signal. Very few to no fibers were detected to be fluorescently labeled in the sciatic nerve of mouse lines CNPMTO-C and F. Finally, teased fiber preparations revealed that the tdTomato protein is specifically enriched in cytoplasm-rich structures of Schwann cells rather than in compact myelin. It is specifically localized at the loosely-packed Schmidt-Lanterman incisures visible as intensely fluorescing stripes in the myelin sheath (Fig. 7 A^t^ 1+2), the nodes of Ranvier (Fig. 7 A^t^ 3+4), and the inner and outer tongues of myelin (Fig. 7A^c^, A^t^ 5+6, black and white arrows). The inner and outer tongues can be identified in longitudinal and cross orientation of both small- and large-diameter myelinated axons.

**Fig. 7:**
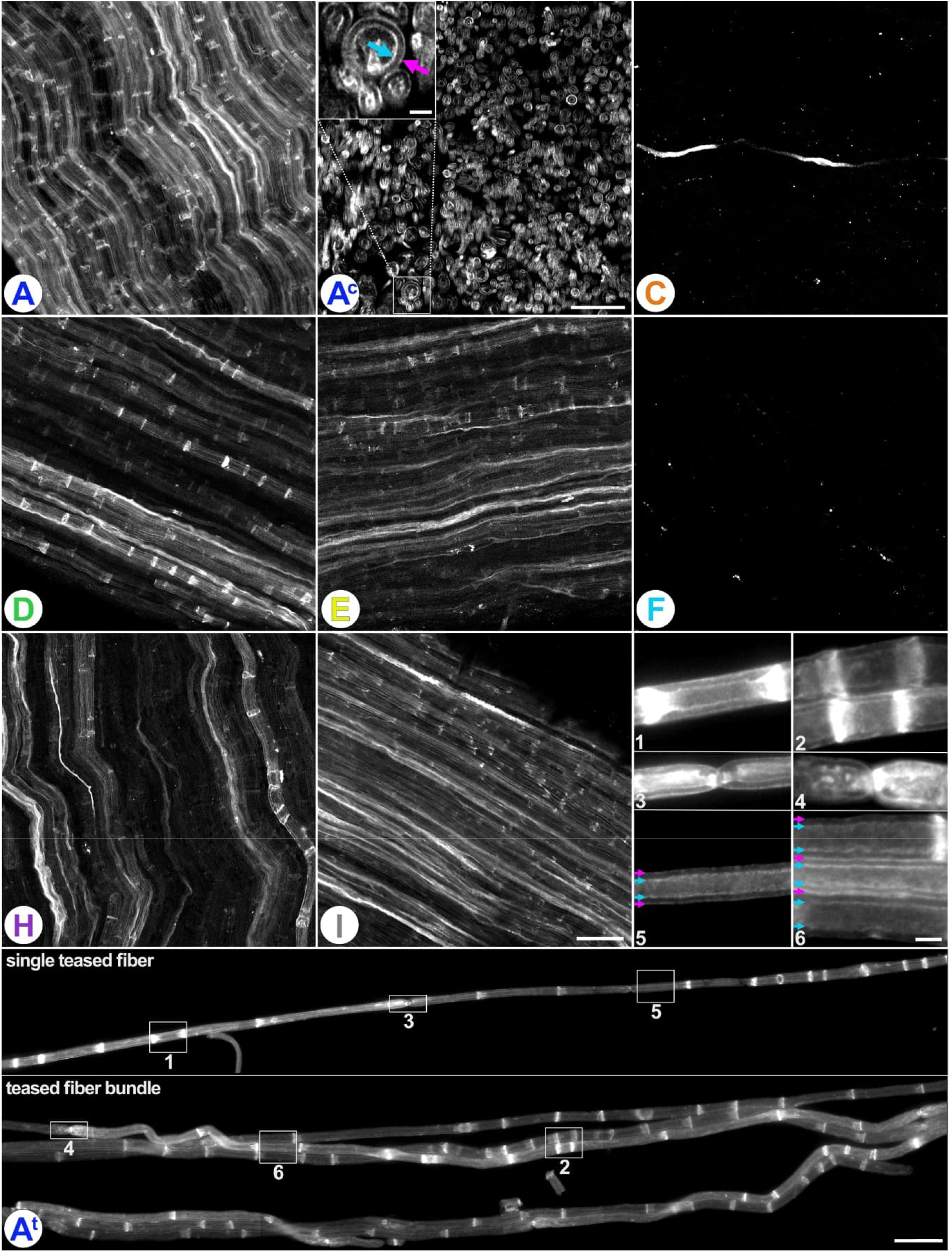
tdTomato expression in sciatic nerve fibers. **A-I:** Whole-mounted sciatic nerves of all mouse lines, the letters in the circles indicate the respective mouse line. Scale bar for CNPMTO-I corresponds to 40 µm and applies to panel A to I. **A**^**c**^: cross-section of the sciatic nerve of line CNPMTO-A. Scale bars 40 µm and 5 µm (inset). **A**^**t**^: teased fibers of the sciatic nerve of line CNPMTO-A. Magnifications 1+2 highlight Schmidt-Lanterman incisures, 3+4 nodes of Ranvier, 5+6 the inner and outer tongues. Scale bars 40 µm (lower panels) and 5 µm (magnifications labeled 1-6). Cyan and purple arrows indicate the inner and outer tongue of myelin, respectively, in panels A^c^ and A^t^ 5+6.

## Discussion

Fluorescence microscopy of cellular structures and functional dynamics has revolutionized the study of the nervous system in both health and disease. Labeling of specific cell types by expression of fluorescent proteins allows for studying their structure and structural dynamics, while cell-type specific expression of functional reporters, including genetically encoded Ca^2+^ or metabolite sensors, enable to assess the dynamics of these cellular processes both *in vitro* and *in vivo* (e.g. Hirrlinger et al., 2004; Nimmerjahn et al., 2005; Madisen et al., 2012; Trevisiol et al., 2017; Zuend et al., 2020; Looser et al., 2024; Thieren et al., 2025). However, many functional sensors are fluorescent within the green/yellow spectral range, thereby hampering their simultaneous visualization together with cellular structures via GFP-based structural reporters including in mouse lines previously used for structural imaging of myelin based on membrane-targeted versions of EGFP (Chong et al., 2012; Deng et al., 2014). To overcome these limitations, we here present a set of seven novel mouse lines which express the red fluorescent, membrane-tethered tdTomato in myelinating glial cells. These lines enable robust, specific, and spectrally distinct labeling of myelin in the red spectral range in both the CNS and PNS, providing a powerful tool for multimodal imaging in myelin research.

Membrane-anchoring of EGFP has been a successful strategy to label myelin both in mice (Chong et al., 2012; Deng et al., 2014) and zebrafish (Almeida et al., 2011; Arafa et al., 2026) taking advantage of the CAAX-membrane anchor motif of human H-Ras which facilitates prenylation of the protein (Hancock et al., 1989). This sequence was also used to localize tdTomato to the plasma membrane of macrophages in zebrafish (Oehlers et al., 2015), suggesting its applicability for tdTomato. Indeed, recent adeno-associated virus (AAV)-mediated strategies have been introduced to target membrane-bound tdTomato to oligodendrocytes for *in vivo* imaging (Snaidero et al., 2020; Mezydlo et al., 2023). However, AAV-based approaches can show variable transduction efficiency across brain regions and experimental conditions, which may limit their suitability for longitudinal studies. In contrast, transgenic expression provides stable and reproducible labeling, facilitating long-term imaging and combination with functional sensors in other cell types or axons. In this study, transgenic expression of CAAX-tagged tdTomato under the control of the *Cnp* promoter resulted in the generation of seven transgenic mouse lines with robust and specific labeling of myelinating glial cells in both the CNS and PNS by the red fluorescence of tdTomato. tdTomato expression is wide-spread across major CNS regions, both in gray and white matter, but also extends to the PNS, as evidenced by labeling of myelin in the sciatic nerve. As commonly observed for many transgenic mouse lines (e.g. Feng et al., 2000; Hirrlinger et al., 2005), the expression patterns differ between the seven CNPMTO mouse lines, which is also reflected by the quantification of tdTomato-labeled myelin sheaths as identified by MBP-immunopositivity, most likely due to the inherent variability in transgene integration sites and copy number. Importantly, this diversity provides a toolkit of complementary lines allowing selection of the respective mouse line based on the specific experimental aims. For example, wide-spread coverage of most myelin sheaths as in line CNPMTO-A will enable experiments addressing myelin loss during pathology, while sparse expression in line CNPMTO-C provides single-cell resolution allowing to delineate individual oligodendrocytes and their associated myelin sheaths and to quantify their internodal lengths.

Myelin sheaths mainly consist of densely packed, compacted myelin membranes. During compaction, which crucially depends on MBP (Snaidero et al., 2017), cytosol, including most cytosolic proteins, is extruded from between the membranes with MBP acting as a molecular sieve that excludes large proteins from compact myelin (Aggarwal et al., 2011; Simons et al., 2024). However, at the inner and outer tongues and the paranodal loops (in the PNS also at Schmidt-Lanterman incisures) myelin is non-compact and thus comprises cytosol in a system also referred to as myelinic channels which is highlighted by soluble, non-membrane anchored tdTomato (Edgar et al., 2021). While in the mouse lines presented here, the entire myelin sheaths are clearly visible, most of the membrane-targeted tdTomato is located along the myelinic channels as evidenced by immuno-transmission electron microscopy in optic nerves, as well as in the sciatic nerve in which Schmidt-Lanterman incisures, paranodal loops, and the inner and outer tongues show the strongest fluorescence. Furthermore, in cross sections of the sciatic nerve two fluorescence rings are observed around each axon, most likely correlating to the inner and outer tongue of myelin. Therefore, tdTomato appears largely excluded from compact myelin, which might be a drawback for experiments in which explicit labeling of compact myelin is desired. Interestingly, a very similar pattern of fluorescence in Schmidt-Lanterman incisures and the inner and outer tongue was observed in a mouse line with membrane-tagged EGFP (Deng et al., 2014), suggesting that such localization is not specific for tdTomato and thus not due to the higher molecular weight of tdTomato (54.2 kDa) compared to EGFP (26.9 kDa). Furthermore, as the *Cnp*-mEGFP mouse line (Deng et al., 2014) has been widely used to study myelin and its dynamics, the similar-appearing localization pattern suggests that the cellular distribution of tdTomato observed in the novel mouse lines presented here does not generally hamper the usability of these mice.

The novel CNPMTO mice reported here offer a combination of features, which were not available as a transgenic tool so far. While previously reported lines expressing cytosolic EGFP under the control of the *Plp1*- (Mallon et al., 2002) or the *Mobp*-promoter (Gensat; Gong et al., 2003; Hughes et al., 2018) or membrane-anchored EGFP driven by the *Mbp*-(Chong et al., 2012) or the *Cnp*-promoter (Deng et al., 2014) enable labeling of myelinating glial cells and their myelin sheaths, their green fluorescence limits a combined use with functional sensors emitting in the green to yellow spectral range. In the *Plp1*-DsRed mouse line, the red fluorescent protein DsRed is expressed in oligodendrocytes, resulting in specific labeling of these cells (Hirrlinger et al., 2005). While the red fluorescent protein would be compatible with green functional sensors, myelin sheaths are only very weakly labeled in this mouse line due to the formation of DsRed aggregates, thereby hampering structural analysis of myelin. Alternatively, genetic labeling of myelin could be generated by combining mouse lines with expression of Cre or CreERT2 in myelinating cells with appropriate reporter mice expressing red fluorescent proteins after Cre-mediated DNA recombination (e.g. Edgar et al., 2021). However, such an approach is not compatible with the simultaneous application of Cre-mediated DNA recombination for conditional gene deletion and requires extensive mouse breeding. Furthermore, when using CreERT2, application of tamoxifen may have unwanted side-effects. Therefore, the novel mouse lines presented here fill a critical gap by direct transgenic expression of a red fluorescent, membrane-targeted, myelin-specific reporter with clearly labeled myelin sheaths and spectral compatibility with green functional sensors.

In conclusion, we present a novel and versatile tool for imaging of myelinating glial cells and their myelin sheaths in *in vivo, ex vivo* and *in vitro* experiments. The CNPMTO mouse lines allow visualization of myelin structure with red fluorescence in both CNS and PNS, providing a platform to investigate structural and functional aspects of myelin *in vivo*, particularly in combination with green fluorescent, genetically encoded functional sensors expressed in neurons or glial cells. These lines hold great promise towards advancing our understanding of myelin structure, dynamics, and plasticity, axoglial metabolic interactions, and the pathophysiology of myelin-related disorders.

## Acknowledgement

We are grateful to the Laboratory for Transgenic Technologies at the Animal Facility of the MPI-NAT City-Campus for their excellent work generating the transgenic mice. We thank Dr. Anke Schraepler, Dr. Sarah Kimmina and the staff of the mouse facility for mouse husbandry and support with the associated bureaucracy as well as Ulli Bode, Boguslawa Sadowski, Christin Kellner and Grit Marx for outstanding technical support. We thank Prof. Markus Morawski for sharing his expertise with advanced microscopy.

## Supplementary material

**Supplementary Figure 1:**
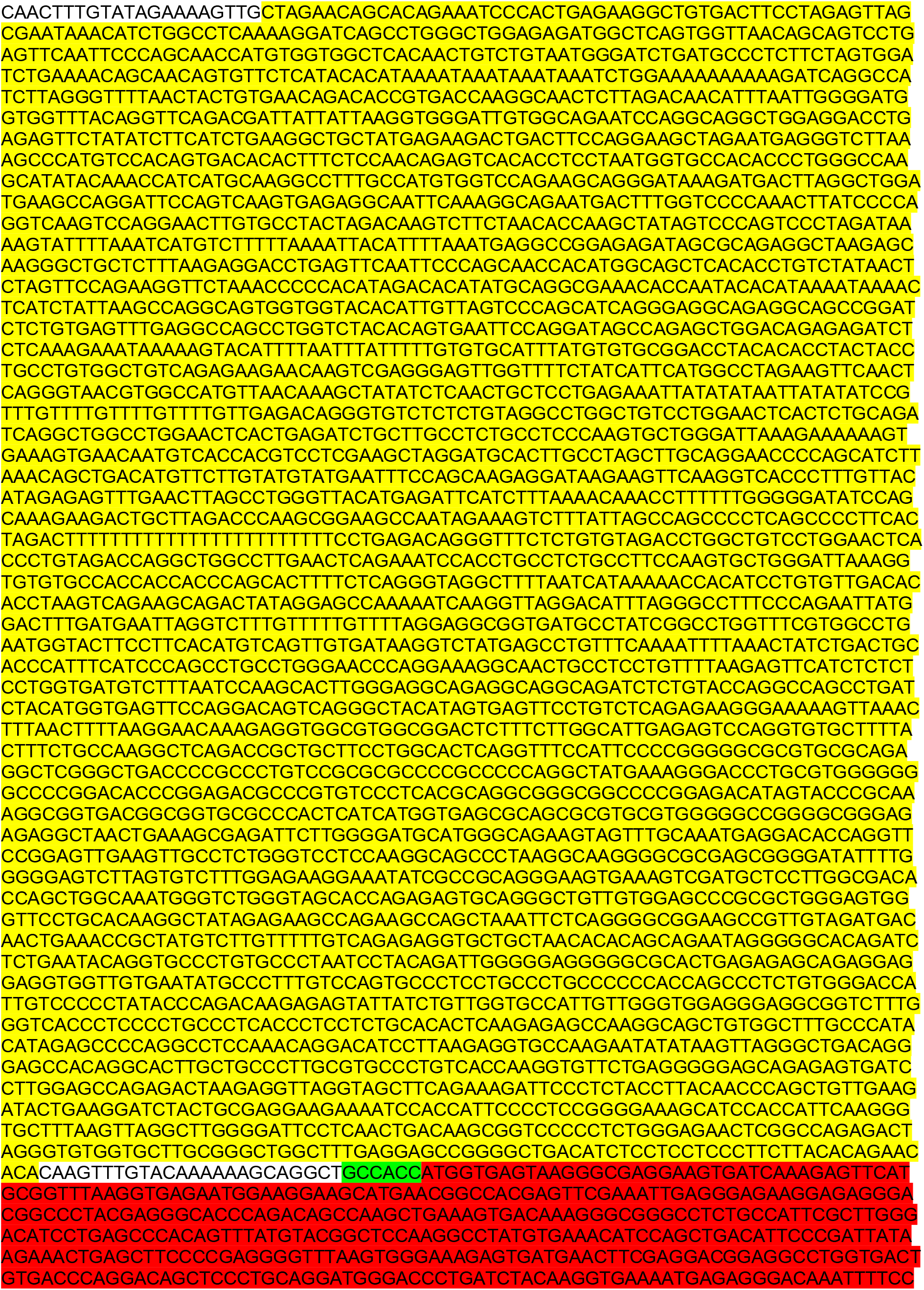
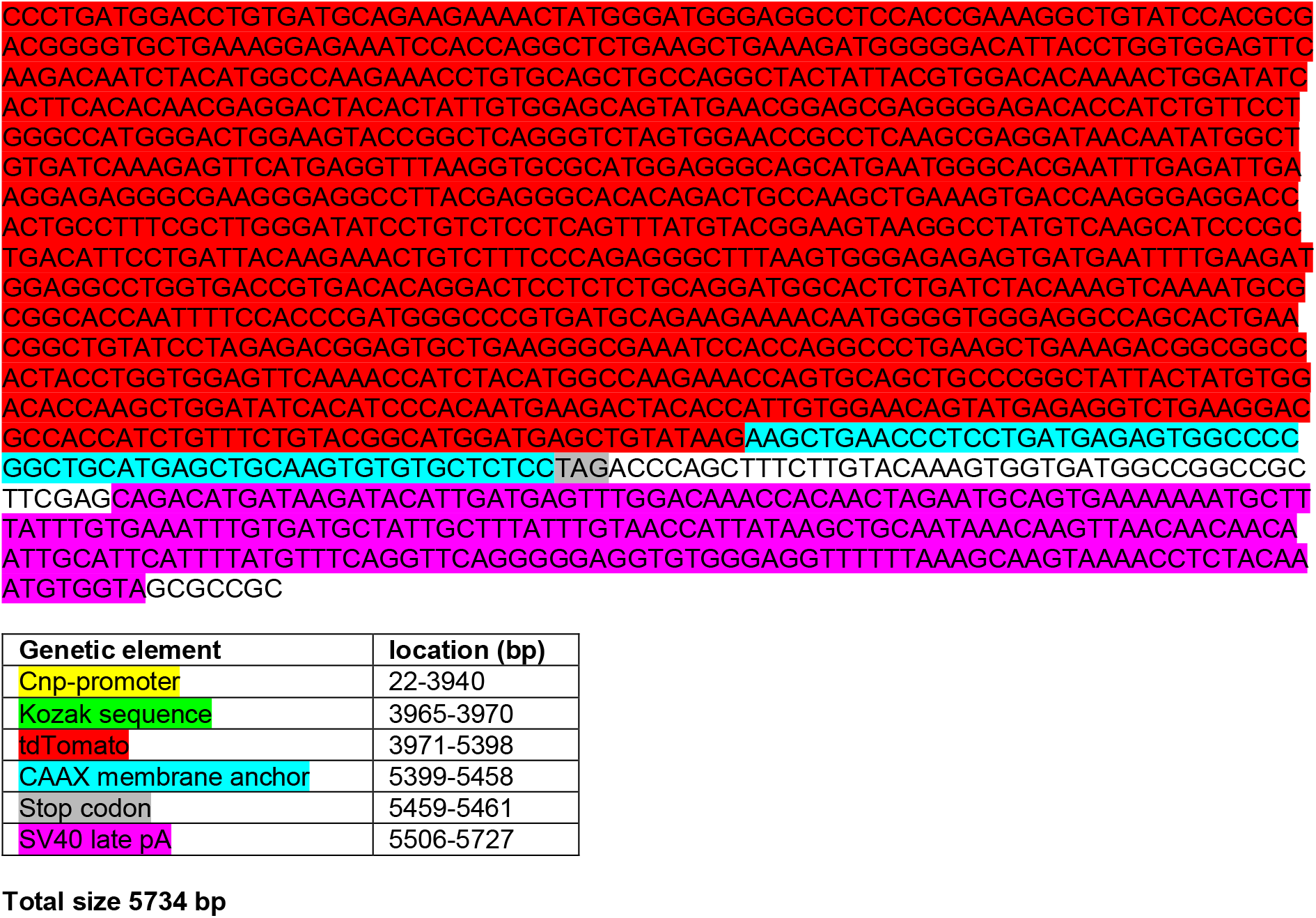
Transgene construct.

